# Evaluating MaxEnt Modeling Strategies for Predicting Suitable Habitats of Invasive Insects Under Climate Change Scenarios

**DOI:** 10.64898/2026.04.18.719331

**Authors:** Pankaj Chouhan, Olmo Zavala-Romero, Muhammad Haseeb

## Abstract

Invasive insect species pose serious threats to agriculture and ecosystems, with their spread increasingly accelerated by global trade and climate change. To support prevention and mitigation efforts, it is essential to map the regions where these pests can survive and thrive. Here, we apply MaxEnt, a leading species distribution modeling framework, to estimate current (2020) and future (2040–2060) suitable habitats for five major invasive insects across the contiguous United States: brown marmorated stink bug, corn earworm, spongy moth, root weevil, and spotted lanternfly. To account for an uncertain climatic future, these projections are generated under four shared socioeconomic pathways, which reflect a range of plausible climate change scenarios.

Beyond forecasting distributions, we examine several key modeling decisions, especially those often overlooked in practice. In particular, we find that background sampling strategies play a critical role in model calibration and that a hybrid sampling approach with a moderate buffer bias provides better predictive accuracy. We also show that permutation importance scores, commonly used to rank environmental variables, are highly sensitive to small changes in the background data and should be interpreted with caution.

Finally, to bridge the gap between ecological modeling and applied machine learning, we provide a self-contained, math-focused background to MaxEnt aimed at practitioners outside of traditional ecological fields. Overall, this work delivers reproducible modeling workflows and critical insights into building robust, transparent, and ecologically meaningful MaxEnt models for climate-informed species distribution analysis.

## 1. Introduction

Invasive insect species pose a critical biological and economic threat. They are highly successful in new habitats because they typically lack the natural predators and specific pathogens that kept their populations in check in their native ranges, allowing them to adapt and proliferate rapidly. These species are a threat to native agriculture, forestry, and ecosystems. Worldwide, the damage caused by invasive species totals over $423 billion each year Bradshaw et al. (2024). For instance, in the US alone, invasive pests caused an estimated $21 billion in annual losses between 2010 and 2020 through crop damage, control efforts, and environmental harm Fantle-Lepczyk et al. (2022). This already severe challenge is now being compounded by accelerating climate change. Rising global temperatures and shifting weather patterns create new, favorable environmental conditions that allow these pests to expand their geographical ranges, survive milder winters, and complete more breeding cycles per season.

To manage these challenges, it is important to understand where pest insects are likely to live and thrive. Mapping their *suitable habitats*, or *ecological niches*, helps us identify the environmental conditions they need to survive. This information is useful for studying their spread, and for designing control strategies. As temperatures and precipitation patterns shift due to climate change, pest insects are moving into new regions, putting ecosystems, agriculture, and food security at greater risk. Predicting where these insects might spread in the future is useful for taking early action, improving surveillance, and reducing their impact.

Species Distribution Models (SDMs) are a suite of models that can estimates suitable habitats using environmental conditions information. These models can then be used to predict the suitable habitats under both current and projected climate scenarios. Among these, maximum entropy (*MaxEnt*) is one of the most popular SDMs. MaxEnt works by comparing environmental conditions at known presence locations (where insects have been observed) with background points (that represent the range of environmental conditions across the entire study area).

MaxEnt offers strong predictive performance and is flexible in handling diverse environmental variables Phillips et al. (2004, 2006); Elith et al. (2011); Merow et al. (2013); Barve et al. (2011). While powerful, MaxEnt’s output is highly sensitive to user decisions such as the selection and configuration of background points, feature types, and regularization settings. These choices affect not only the MaxEnt performance but also the ecological interpretation and transferability of its predictions (Elith et al., 2011; Phillips et al., 2017).

In this paper, we adopted a practical, data-driven workflow for using MaxEnt to model the distribution of five major invasive insect species in the contiguous US: brown marmorated stink bug (BMSB), corn earworm (CE), spongy moth (SM), root weevil (RW), and spotted lanternfly (SL). We estimate suitable habitats for the species for the present (2020) and future time points (2040, and 2060). These projections are modeled under four climate change scenarios, known as shared socioeconomic pathways (SSPs). SSP1–2.6 represents a low-emission, sustainable future, while SSP2–4.5 assumes moderate development and emissions. SSP3–7.0 reflects a fragmented world with high emissions, and SSP5–8.5 models a fossil fuel–driven future with very high warming.

While much of the existing literature focuses on ecological framing, we present a onestop resource tailored for applied machine learning practitioners, laying down the complete mathematical framework, and helping them navigate MaxEnt’s complexities using clear, reproducible steps.

Beyond predicting suitable habitats, we also critically evaluate how to interpret the permutation importance scores generated by MaxEnt. While commonly used as a way to rank predictor variables, we argue that these metrics can be misleading if taken at face value. Our initial goal was to identify the most influential variables driving habitat changes in response to climate change. However, we found that MaxEnt is better suited to highlight a set of important features rather than to establish a definitive ranking.

## 2. Materials and Methods

### 2.1. Theoretical background and MaxEnt framework

Species distribution models (SDMs) are typically constructed from ecological observations in which species occurrences are recorded as presence locations. A central challenge in this setting is the absence of reliable negative labels: non-detection does not imply true absence due to imperfect sampling and detection processes. As a result, presence-only modeling frameworks are widely adopted, where inference is based solely on observed presences and the environmental conditions across the study region.

In this setting, the objective is to estimate a spatial distribution that reflects habitat suitability as a function of environmental covariates. Let *X* denote the set of candidate locations and *x*_1_*, . . . , x_m_* ⊂ *X* the observed presence sites. Each location *x* is associated with features ***f*** (*x*) ∈ **ℝ***^J^* . Following Phillips et al. (2004), we define *π*(*x*) = *P* (*x* | *y* = 1) as the sampling distribution of observed presences. Under the assumption of uniform sampling over X, Bayes’ rule yields:

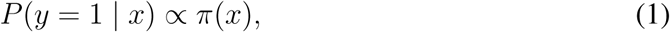

where the proportionality constant depends on the (unknown) species prevalence. Consequently, estimating *π*(*x*) is sufficient to recover relative habitat suitability.

#### 2.1.1. MaxEnt

MaxEnt estimates *π*(*x*) by selecting the maximum-entropy distribution subject to constraints that match empirical feature averages at presence locations. The resulting model belongs to the exponential family:

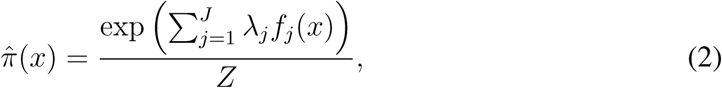

where *Z* is the normalization constant and ***λ*** are feature weights.

The parameters are estimated by maximizing the regularized log-likelihood over presence samples:

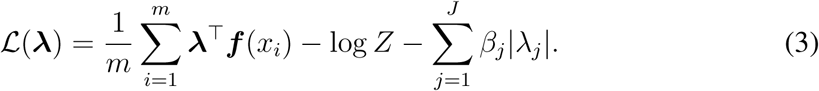

Regularization is critical to control overfitting, particularly when the feature space is expanded. In practice, MaxEnt augments the original covariates using transformations such as linear, quadratic, product, hinge, and threshold features, enabling flexible nonlinear responses. To balance contributions across features, regularization terms are scaled according to feature variability:

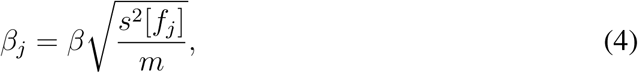

where *s*^2^[*f_j_*] denotes the empirical variance of feature *f_j_*.

*Logistic output..* The raw MaxEnt output *π*^(*x*) depends on the size of the background sample and is not directly interpretable. To obtain a calibrated measure of presence likelihood, MaxEnt can be expressed as a joint maximum-entropy model over (*x, y*), leading to the conditional probability:

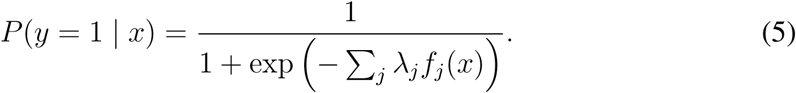

This logistic transformation provides an interpretable measure of habitat suitability and establishes a formal connection between MaxEnt and logistic regression.

### 2.2. Data

#### 2.2.1. Species occurrence data

In this study, we considered invasive pest insects that pose a threat to US agriculture and native ecosystems. Specifically, we considered the five insects: brown marmorated stink bug, corn earworm, spongy moth, root weevil, and spotted lanternfly. These insects were artificially introduced into the US via means of trade or by human error Rice et al. (2014); Lapointe et al. (2014); Urban and Leach (2023). Since their introduction, they have rapidly spread and caused significant damage because of the absence of natural predators. A brief explanation of the characteristics of each insect is presented in the supplementary information (SI).

Species occurrence data for these species were obtained from publicly available records provided by the Global Biodiversity Information Facility (GBIF). Figure 1 illustrates the current distribution of these insects, along with the number of observations available for each species in the contiguous US.

**Figure 1:**
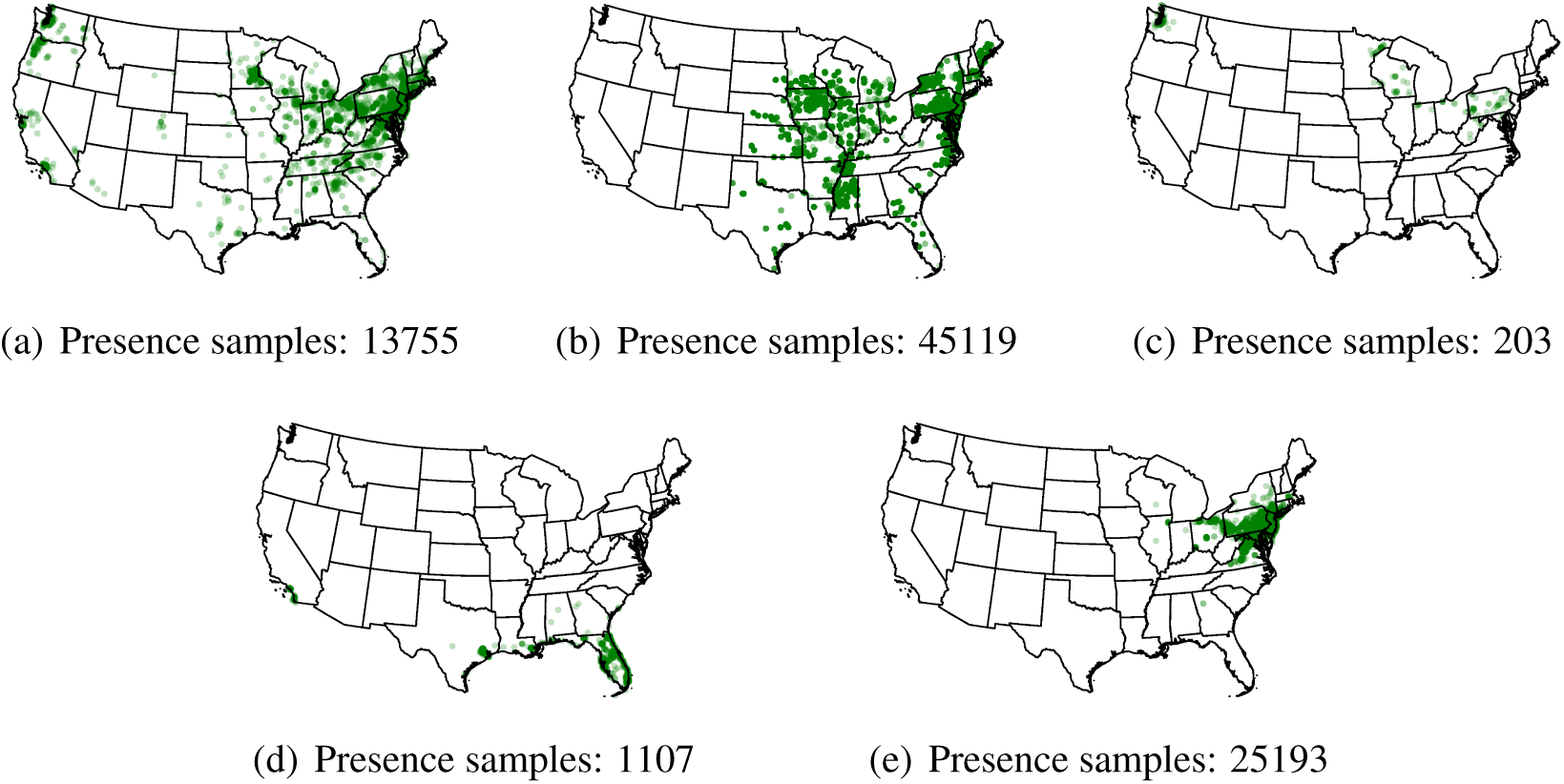
Distribution of presence samples across the contiguous US for (a) brown marmorated stink bug, (b) corn earworm, (c) spongy moth, (d) root weevil, and (e) spotted lanternfly.

#### 2.2.2. Environmental predictors

We use a set of 19 bioclimatic variables derived from temperature and precipitation data, available at a spatial resolution of 10 arc-minutes. To train the model, we use the average climate data from the period 1970–2000. Future projections are based on multi-decadal averages corresponding to the periods 2021–2040 and 2041–2060. We consider emission scenarios: SSP1-2.6, SSP2-4.5, SSP3-7.0, and SSP5-8.5. All these data are publicly available at the WorldClim website. Apart from these, we also consider the elevation profile of the US as a variable in our study.

### 2.3. Modeling pipeline and evaluation

Following best practices in the MaxEnt modeling literature, we developed a preprocessing pipeline tailored to the needs of our study. The first step addresses a common issue with presence-only data: spatial bias. Such data are often recorded in areas with high human accessibility, like near roads or rivers Elith et al. (2011); Phillips et al. (2017). If left uncorrected, this bias can distort the species’ estimated ecological niche and reduce the model’s ability to accurately identify suitable habitats Iturbide et al. (2018); Grimmett et al. (2020); Liu et al. (2019); Barbet-Massin et al. (2012).

To mitigate this bias, we applied spatial thinning Castellanos et al. (2019); Elith et al. (2011), using a thinning distance of 10 kilometers (km)—roughly matching the spatial resolution of our bioclimatic variables. This choice balances the trade-off between reducing bias and retaining sufficient data for model training: smaller distances may not sufficiently address clustering, while larger ones may discard too many records, weakening model performance. Table 1 summarizes the number of presence points before and after thinning.

**Table 1:** Number of presence points before and after spatial thinning.

|  | Original | Spatially thinned |
| --- | --- | --- |
| BMSB | 13,755 | 1412 |
| CE | 45,119 | 719 |
| SM | 203 | 193 |
| RW | 1,107 | 185 |
| SL | 25,193 | 664 |

We also addressed multicollinearity among bioclimatic variables by removing pairs with a Pearson correlation coefficient ≥ 0.9. This improves model interpretability and reduces the risk of overfitting.

Another key aspect of MaxEnt modeling is the configurations of background points. Decisions around their spatial extent, density, and distribution can significantly affect predicted probabilities Merow et al. (2013); Phillips et al. (2017); Whitford et al. (2024). For the spatial extent of the study, we followed the recommended guidelines of distributing background points across the full range of environmental conditions in the contiguous US Merow et al. (2013). Regarding the number of points, previous studies have recommended using either a fixed multiple of the number of presences or a standardized total (e.g., 10,000) Fourcade et al. (2014); Bennett et al. (2019); Elith et al. (2011); Barber et al. (2022). Our preliminary experiments supported the latter strategy, and we therefore fixed the number of background points at 10,000 for all models.

To select the best distribution of background points, we performed a series of experiments (Sec. 3.1) that studied the effect of background point distribution against the presence point distribution. We employed a mixed sampling strategy Shipley et al. (2022), which integrates uniform and buffer-based background point sampling through a mixing coefficient *m*. This method enables a tunable balance between global and local environmental contrasts in the training data. When *m* = 0, all background points are sampled uniformly; when *m* = 1, all are drawn from the buffer. Intermediate values of *m* produce a controlled blend of the two. This can be seen in Fig. 2. Our results (Sec. 3.1) indicate that selecting an intermediate value of *m* offers a practical compromise, helping to reduce bias while maintaining generalizability. This finding aligns with results from Whitford et al. Whitford et al. (2024), who also observed performance gains using a mixed background sampling strategy.

**Figure 2:**
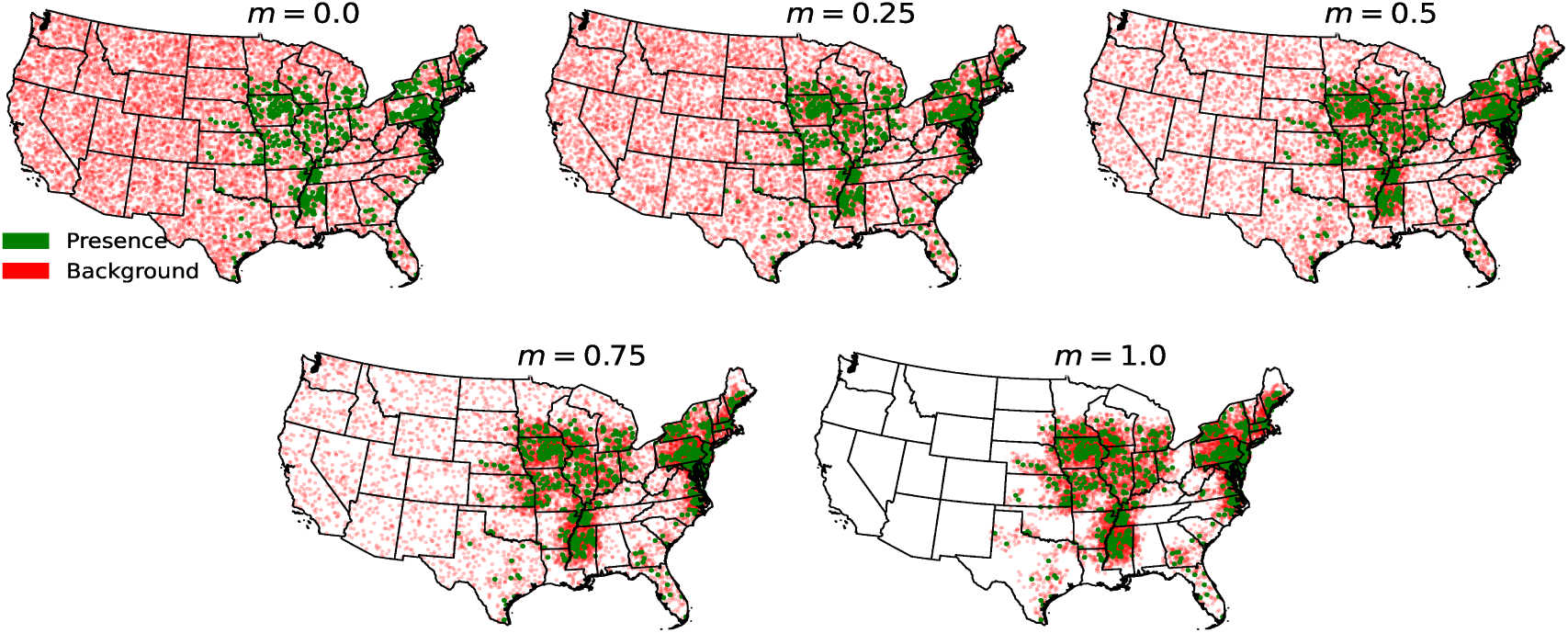
Distribution of background points for the corn earworm experiment with different values of the mixing coefficient *m*. A value of *m* = *x* indicates that *x*% of the background points are sampled from a buffer around the presence locations, while the remaining (100 − *x*)% are drawn uniformly from the study area.

To create a train-test split for the presence samples, we used a checkerboard partitioning strategy. This approach ensures that each spatial region contributes to both the training and test sets, promoting a roughly balanced split and further reducing spatial bias Muscarella et al. (2014). Figure 3 illustrates this method for the corn earworm experiment. For test-set, the background point are sampled uniformly. This ensures that the model is evaluated on its ability to extrapolate.

**Figure 3:**
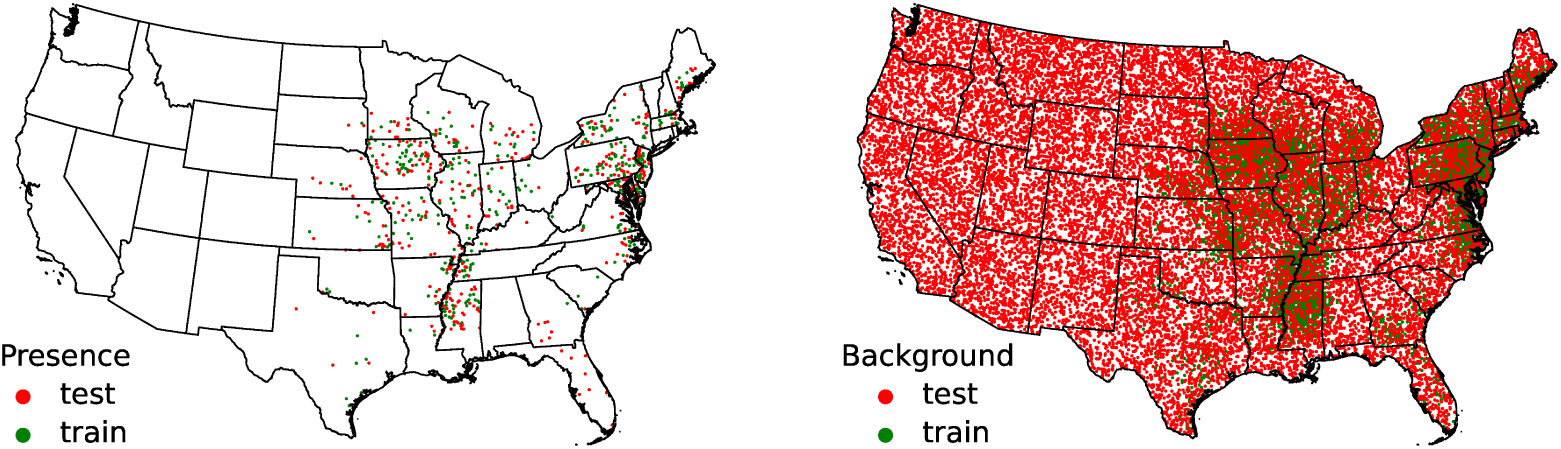
The distribution of presence and background points for the train-test split.

Finally, we computed two metrics to evaluate our model: area under the curve (AUC) and the continuous Boyce index (CBI). AUC is inappropriate for presence-only data, as it is most effective when true absence data are available, such as in classification problems Merow et al. (2013); Lobo et al. (2008). It is also sensitive to the geographic extent of the study area Merow et al. (2013); Lobo et al. (2008). In contrast, CBI is specifically designed for presence-only data, as it evaluates the model’s ability to distinguish observed presences from a random distribution without relying on pseudo-absences Hirzel et al. (2006). While we report both metrics, all model evaluations are based on the CBI.

To identify the optimal parameter settings, we performed a grid search using the following combinations of feature types: {L, LQ, LQH, LPH, LQHP, LQHPT}, and regularization multipliers *β_j_*∈ {0.5, 1.0, 1.5, 2.0, 2.5}.

## 3. Results

### 3.1. Effect of background point distribution

Figures 4 and 5 illustrate predicted suitable habitats for corn earworm and spongy moth under varying values of the background sampling parameter *m*. As *m* increases from 0 to 1, the fitted models shift from conservative to more permissive prediction.

**Figure 4:**
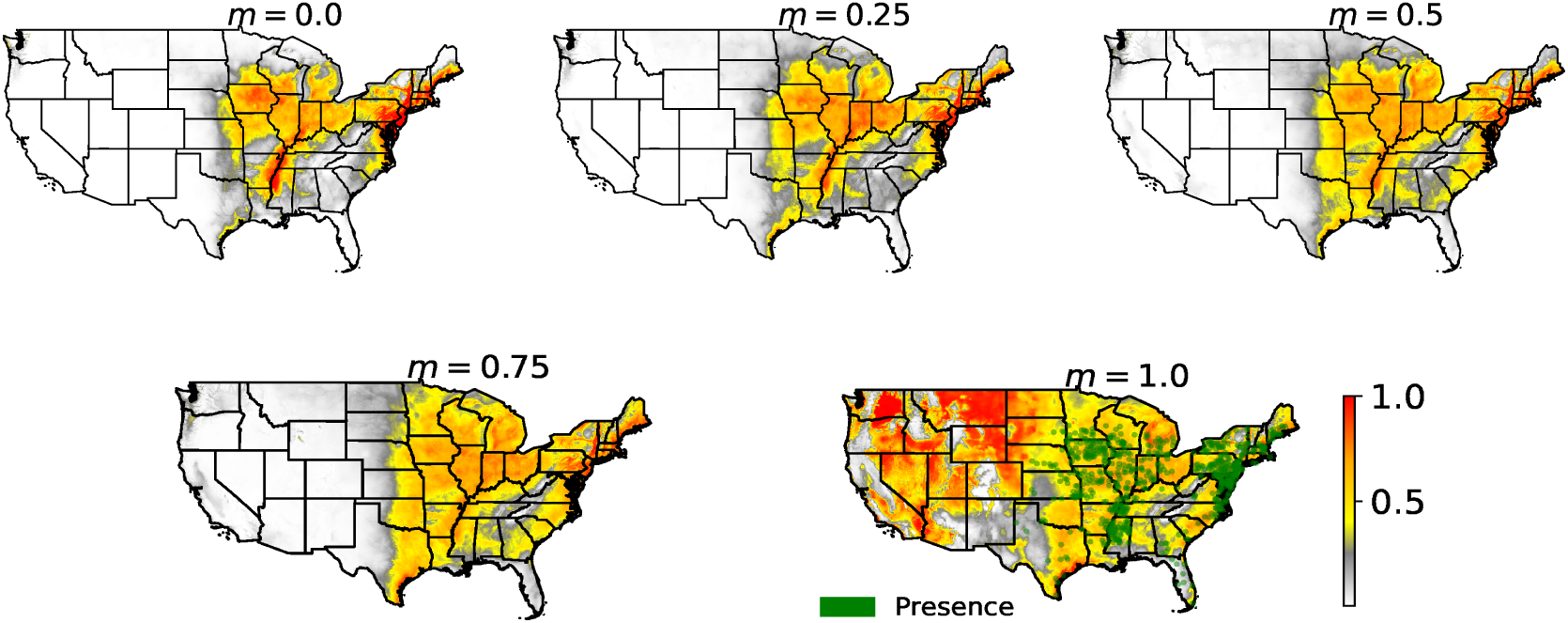
Predicted suitable habitat for the corn earworm using different values of *m*. The model was trained on multi-decadal averages of bioclimatic variables from the 1970–2000 period. The distribution of presence points is shown in the final subplots.

**Figure 5:**
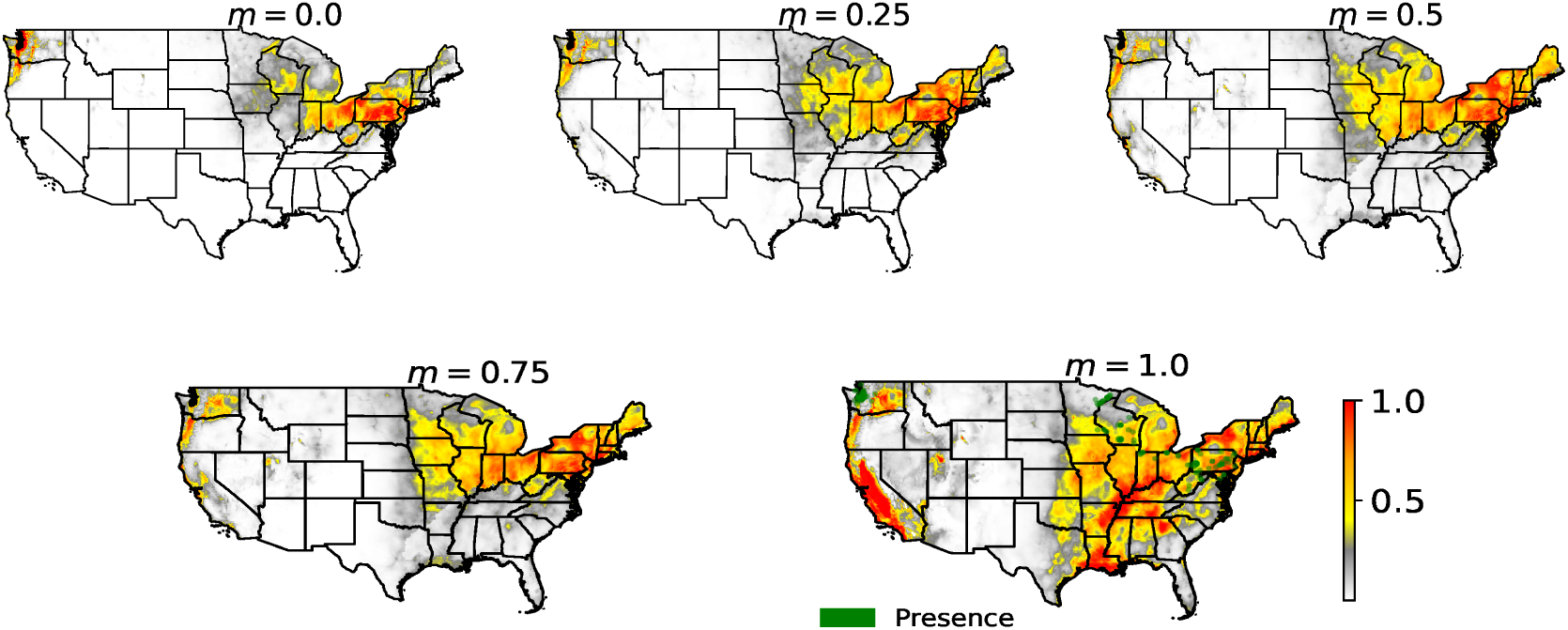
Predicted suitable habitat for the spongy moth using different values of *m*. The model was trained on multi-decadal averages of bioclimatic variables from the 1970–2000 period. The distribution of presence points is shown in the final subplots.

At *m* = 0, background points are sampled uniformly across the study area, producing greater contrast with the presence locations. This results in stronger feature constraints during optimization, forcing the model to deviate significantly from uniformity. Consequently, high suitability is assigned only to regions with environmental conditions closely matching the presence sites, while predictions remain low elsewhere—even in geographically nearby but environmentally distinct areas. For instance, the model predicts low suitability in Georgia (Fig. 4) and Minnesota (Fig. 5) despite presence records, likely because these regions are not well represented in the uniform background.

In contrast, at *m* = 1, background points are drawn from buffers around presence sites, making the environmental background more similar to the presences. This weakens the feature constraints, allowing the model to stay closer to uniform and to generalize more broadly. While this may improve coverage near known occurrences, it can also lead to overprediction in environmentally distinct or unsampled areas.

To quantify which setting of *m* balances these trade-offs, we report the CBI and AUC in Table 2. Across our five species, a modest buffer bias—specifically *m* ≈ 0.25—yields the highest and most stable CBI scores. Four of the five taxa (stink bug, spongy moth, root weevil, lanternfly) peak at *m* = 0.25, and even for corn earworm the gain from *m* = 0.25 to *m* = 0.50 is marginal compared to the dramatic collapse at *m* = 1. By contrast, AUC remains uniformly high (0.88–0.99) for all *m <* 1, even when CBI indicates very poor calibration (e.g., spongy moth at *m* = 0: CBI=0.25 vs. AUC=0.93), and only falls off when CBI is already unacceptable. Moreover, CBI’s standard deviation is low (*<* 0.05) up to *m* = 0.50, but spikes at *m* = 1, reflecting unstable, over-generalized models.

**Table 2:**
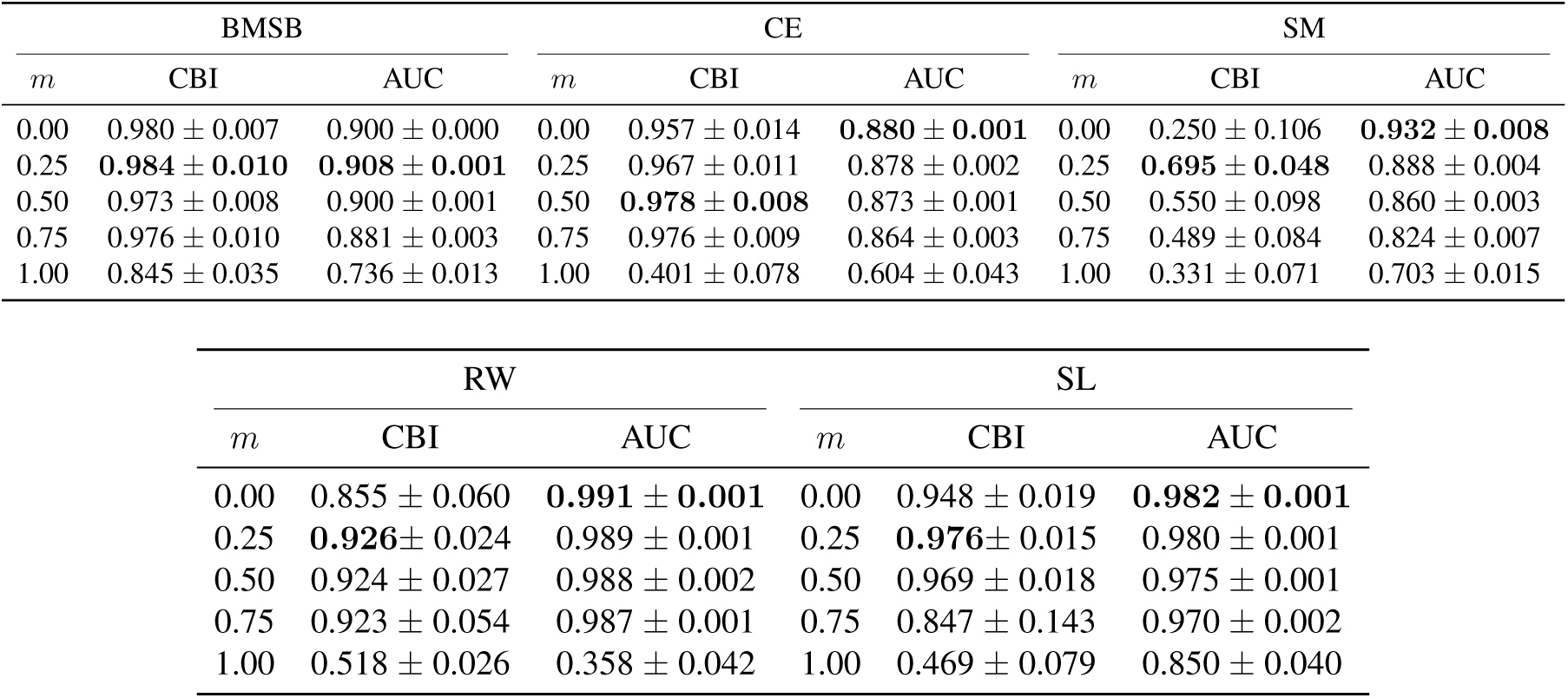
Mean and standard deviation of performance metrics (CBI and AUC) for different mixing coefficients *m*. Each experiment is repeated 20 times. The highlighted value indicates the best observed performance.

| BMSB |  |  | CE |  |  | SM |  |  |
| --- | --- | --- | --- | --- | --- | --- | --- | --- |
| $m$ | CBI | AUC | $m$ | CBI | AUC | $m$ | CBI | AUC |
| 0.00 | 0.980 ± 0.007 | 0.900 ± 0.000 | 0.00 | 0.957 ± 0.014 | **0.880 ± 0.001** | 0.00 | 0.250 ± 0.106 | **0.932 ± 0.008** |
| 0.25 | **0.984 ± 0.010** | **0.908 ± 0.001** | 0.25 | 0.967 ± 0.011 | 0.878 ± 0.002 | 0.25 | **0.695 ± 0.048** | 0.888 ± 0.004 |
| 0.50 | 0.973 ± 0.008 | 0.900 ± 0.001 | 0.50 | **0.978 ± 0.008** | 0.873 ± 0.001 | 0.50 | 0.550 ± 0.098 | 0.860 ± 0.003 |
| 0.75 | 0.976 ± 0.010 | 0.881 ± 0.003 | 0.75 | 0.976 ± 0.009 | 0.864 ± 0.003 | 0.75 | 0.489 ± 0.084 | 0.824 ± 0.007 |
| 1.00 | 0.845 ± 0.035 | 0.736 ± 0.013 | 1.00 | 0.401 ± 0.078 | 0.604 ± 0.043 | 1.00 | 0.331 ± 0.071 | 0.703 ± 0.015 |

| RW |  |  | SL |  |  |
| --- | --- | --- | --- | --- | --- |
| $m$ | CBI | AUC | $m$ | CBI | AUC |
| 0.00 | 0.855 ± 0.060 | **0.991 ± 0.001** | 0.00 | 0.948 ± 0.019 | **0.982 ± 0.001** |
| 0.25 | **0.926 ± 0.024** | 0.989 ± 0.001 | 0.25 | **0.976 ± 0.015** | 0.980 ± 0.001 |
| 0.50 | 0.924 ± 0.027 | 0.988 ± 0.002 | 0.50 | 0.969 ± 0.018 | 0.975 ± 0.001 |
| 0.75 | 0.923 ± 0.054 | 0.987 ± 0.001 | 0.75 | 0.847 ± 0.143 | 0.970 ± 0.002 |
| 1.00 | 0.518 ± 0.026 | 0.358 ± 0.042 | 1.00 | 0.469 ± 0.079 | 0.850 ± 0.040 |

These results suggest treating *m* as a regularization knob in MaxEnt—with low values enforcing strict, conservative predictions and high values leading to over-generalization—and recommend *m* ≈ 0.25 as a default starting point. We further caution against relying solely on AUC for model selection, arguing that calibration-focused metrics like CBI are essential, and note that practitioners should still inspect species-specific performance curves to fine-tune *m*.

### 3.2. Potential distribution under current and future climate scenarios

Using *m* = 0.25, we analyzed the spatial contraction or expansion of suitable habitat area with different MaxEnt settings. We study this across four climate scenarios (SSP1–2.6, SSP2–4.5, SSP3–7.0, and SSP5–8.5) at three time points: current day, 2040, and 2060. For this analysis, we defined suitability as locations where the predicted probability exceeds 0.5—a subjective but interpretable threshold for potential species survival.

Overall, we found that the current predicted habitat aligns well with known population distributions for all insect species, regardless of the MaxEnt model settings. However, projections for future scenarios were sensitive to model configuration. Depending on the chosen settings, we observed both increases and decreases in suitable habitat, with no consistent pattern linking specific configurations to expansion or contraction. Table 3 summarizes the best-performing settings—based on the CBI—for brown marmorated stink bug, root weevil, and spongy moth across different climate pathways.

**Table 3:** Best model configurations based on CBI score under different MaxEnt settings. All experiments were repeated 20 times to ensure statistical robustness, and the habitat expansion column represents the positive or negative change in mean suitable area in the year 2060 relative to the present.

| species | pathway | feature type | $\beta_j$ | habitat expansion |
| --- | --- | --- | --- | --- |
| BMSB | SSP1–2.6 | LQ | 1.5 | True |
| BMSB | SSP2–4.5 | LQH | 1.5 | False |
| BMSB | SSP3–7.0 | LPH | 1.0 | True |
| BMSB | SSP5–8.5 | LQHP | 2.0 | True |
| RW | SSP1–2.6 | LQHP | 2.0 | True |
| RW | SSP2–4.5 | LQHP | 1.0 | True |
| RW | SSP3–7.0 | LPH | 2.5 | True |
| RW | SSP5–8.5 | LQHP | 0.5 | True |
| SM | SSP1–2.6 | LQH | 1.0 | False |
| SM | SSP2–4.5 | LQHP | 1.0 | False |
| SM | SSP3–7.0 | LQH | 0.5 | False |
| SM | SSP5–8.5 | LQHP | 1.5 | False |

We view this inconsistency between model settings and varying future projections as a fundamental characteristic of the MaxEnt modeling framework that is often overlooked. Many studies rely on a single configuration—typically the default settings—without evaluating alternative parameterizations Morales et al. (2017). Our results highlight the limitations of this approach and emphasize the need for more cautious interpretation. The sensitivity of MaxEnt predictions to model settings has been well-documented in prior research Morales et al. (2017); Merow et al. (2013).

To enable comparison with other studies, we provide detailed patterns of habitat expansion and contraction in tables 4–7 (covering all future climate projections), and visualize them in Fig. 6–10 for SSP2–4.5, using the default MaxEnt settings (feature type: LPH, *β_j_*= 1.5). With default settings, we noticed a declining trend in suitable habitat for all species, with the exception of the root weevil, which showed a slight expansion.

**Figure 6:**
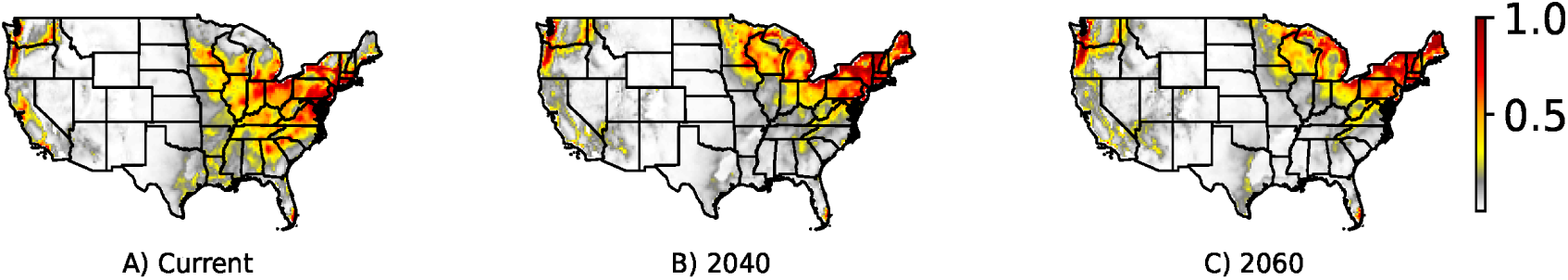
Predicted current and future habitat suitability for the brown marmorated stink bug under SSP2–4.5.

**Figure 7:**
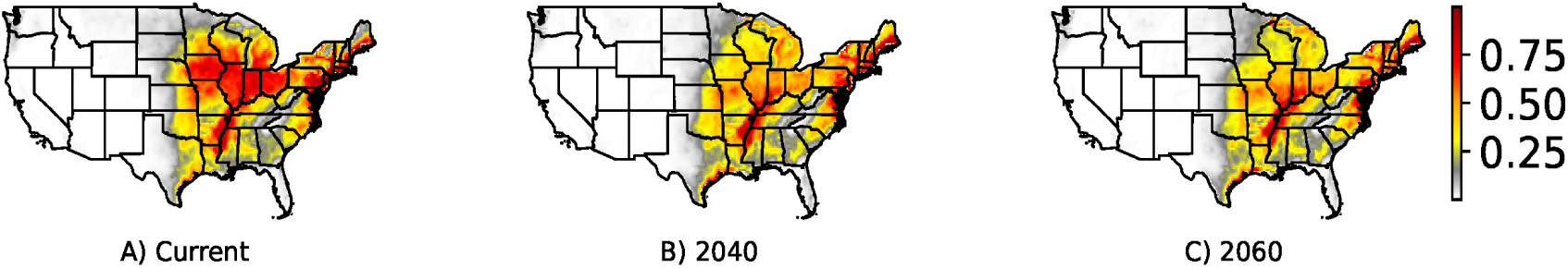
Predicted current and future habitat suitability for the corn earworm under SSP2–4.5.

**Figure 8:**
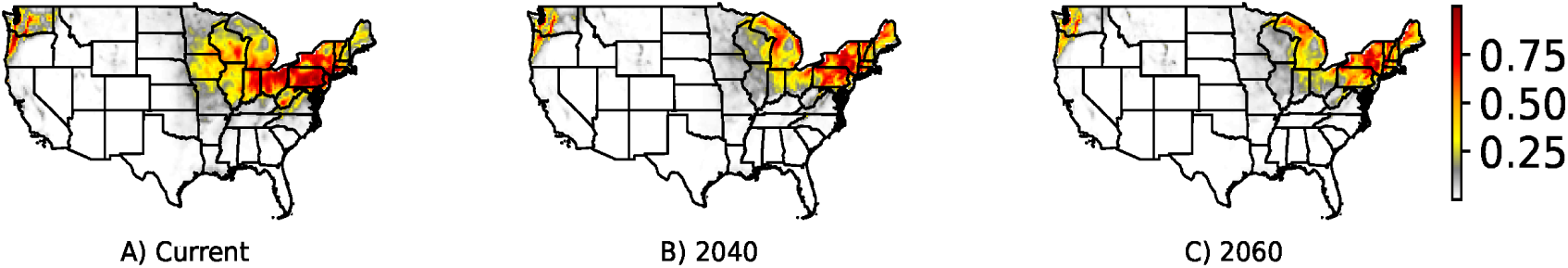
Predicted current and future habitat suitability for the spongy moth under SSP2–4.5.

**Figure 9:**
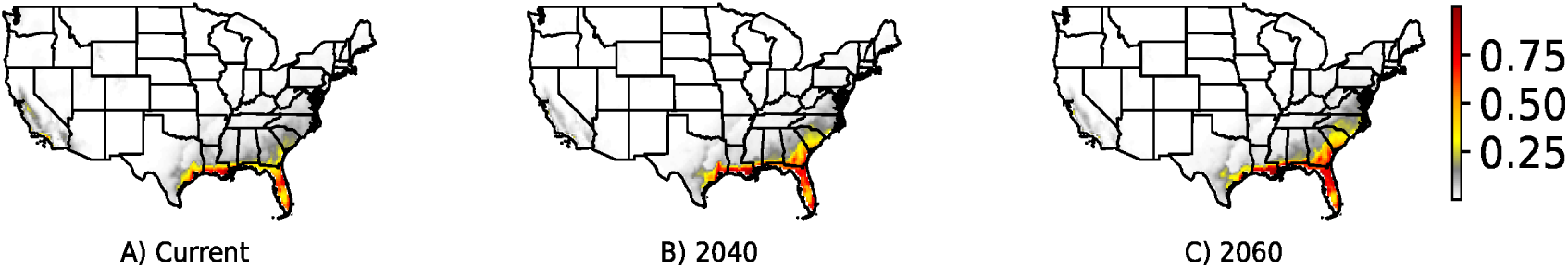
Predicted current and future habitat suitability for the root weevil under SSP2–4.5.

**Figure 10:**
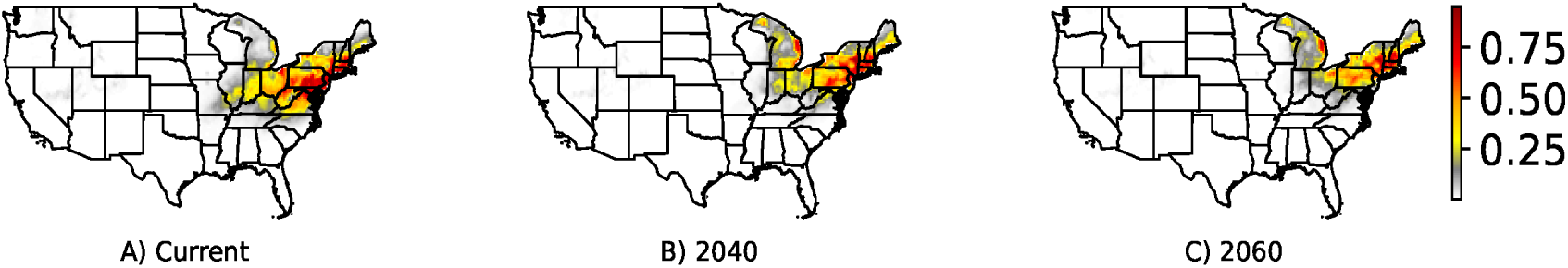
Predicted current and future habitat suitability for the spotted lanternfly under SSP2–4.5.

**Table 4:** Predicted habitat-suitable coverage (fraction of area, %) and percentage change relative to current (mean ± std, %) for five insect species at different time points under SSP1–2.6.

| BMSB |  |  | CE |  |  | SM |  |  |
| --- | --- | --- | --- | --- | --- | --- | --- | --- |
| year | coverage (%) | change (%) | year | coverage (%) | change (%) | year | coverage (%) | change (%) |
| current | 4.29 ± 0.13 | – | current | 6.65 ± 0.15 | – | current | 3.27 ± 0.08 | – |
| 2040 | 3.84 ± 0.35 | –10.5 ± 8.2 | 2040 | 4.60 ± 0.72 | –30.8 ± 10.8 | 2040 | 2.37 ± 0.08 | –27.5 ± 2.4 |
| 2060 | 3.11 ± 0.29 | –27.5 ± 6.8 | 2060 | 4.05 ± 0.98 | –39.1 ± 14.7 | 2060 | 2.23 ± 0.09 | –31.8 ± 2.8 |

| RW |  |  | SL |  |  |
| --- | --- | --- | --- | --- | --- |
| year | coverage (%) | change (%) | year | coverage (%) | change (%) |
| current | 0.73 ± 0.02 | – | current | 1.25 ± 0.06 | – |
| 2040 | 1.17 ± 0.12 | 60.3 ± 16.4 | 2040 | 0.73 ± 0.06 | –41.6 ± 4.8 |
| 2060 | 1.23 ± 0.11 | 68.5 ± 15.1 | 2060 | 0.77 ± 0.07 | –38.4 ± 5.6 |

**Table 5:** Predicted habitat-suitable coverage (fraction of area, %) and percentage change relative to current (mean ± std, %) for five insect species at different time points under SSP2–4.5.

| BMSB |  |  | CE |  |  | SM |  |  |
| --- | --- | --- | --- | --- | --- | --- | --- | --- |
| year | coverage (%) | change (%) | year | coverage (%) | change (%) | year | coverage (%) | change (%) |
| current | 4.12 ± 0.11 | – | current | 6.47 ± 0.15 | – | current | 3.25 ± 0.09 | – |
| 2040 | 4.06 ± 0.37 | –1.5 ± 8.3 | 2040 | 3.10 ± 0.68 | –52.1 ± 11.4 | 2040 | 2.26 ± 0.07 | –30.5 ± 3.1 |
| 2060 | 3.69 ± 0.29 | –10.4 ± 6.8 | 2060 | 2.20 ± 0.99 | –66.0 ± 16.1 | 2060 | 1.98 ± 0.10 | –39.1 ± 4.0 |

| RW |  |  | SL |  |  |
| --- | --- | --- | --- | --- | --- |
| year | coverage (%) | change (%) | year | coverage (%) | change (%) |
| current | 0.72 ± 0.02 | – | current | 1.26 ± 0.06 | – |
| 2040 | 1.24 ± 0.09 | 72.2 ± 12.5 | 2040 | 0.69 ± 0.10 | –45.2 ± 7.1 |
| 2060 | 1.29 ± 0.13 | 79.2 ± 18.1 | 2060 | 0.43 ± 0.11 | –65.9 ± 8.7 |

**Table 6:**
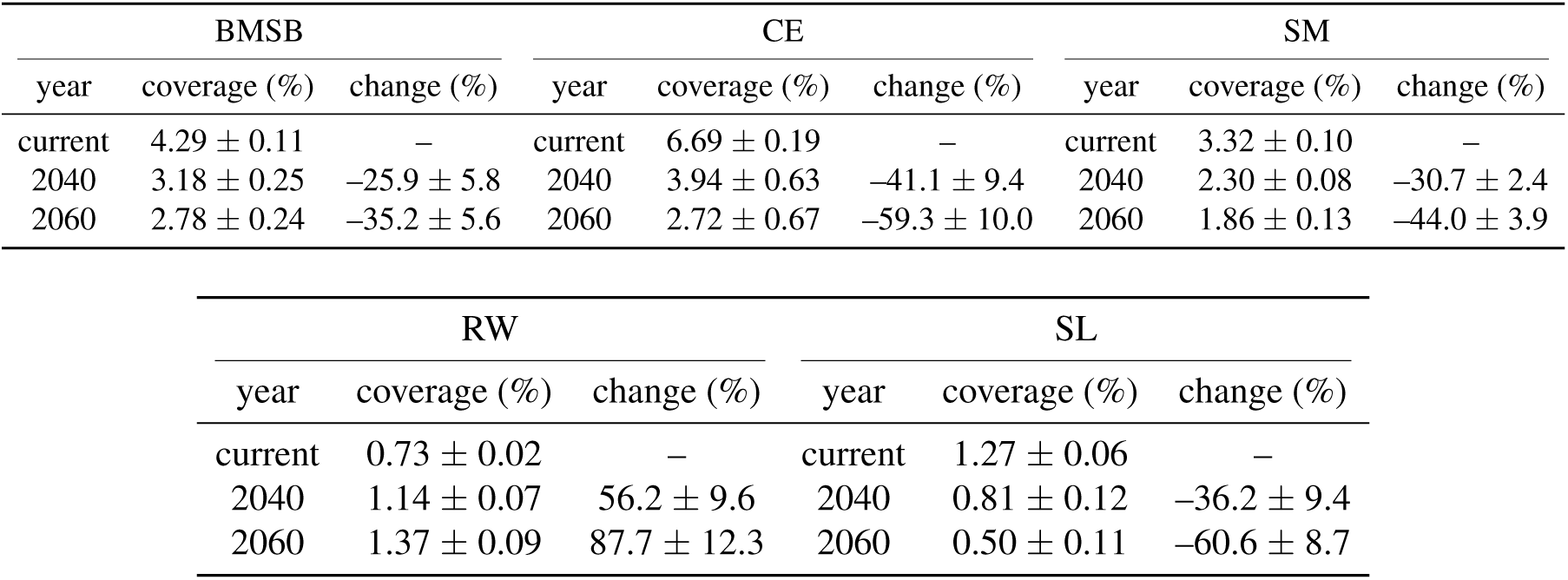
Predicted habitat-suitable coverage (fraction of area, %) and percentage change relative to current (mean ± std, %) for five insect species at different time points under SSP3–7.0.

**Table 7:**
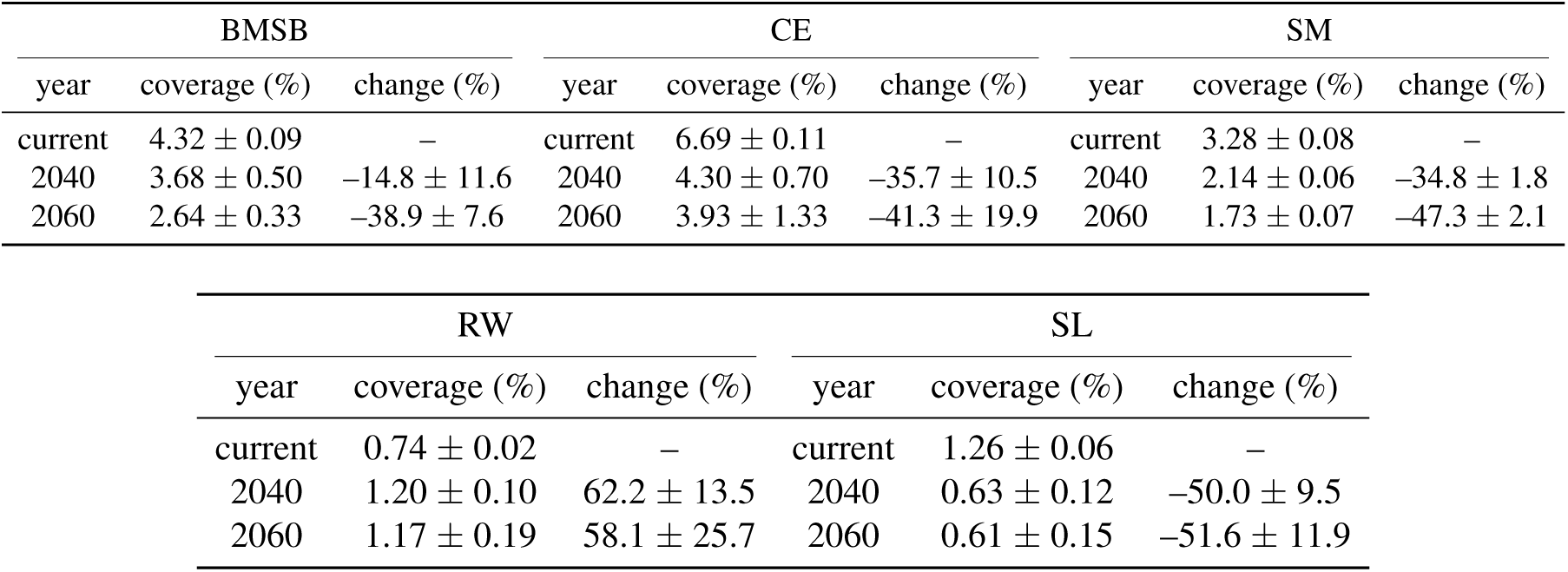
Predicted habitat-suitable coverage (fraction of area, %) and percentage change relative to current (mean ± std, %) for five insect species at different time points under SSP5–8.5.

By 2040, most species show modest losses in suitable habitat. The brown marmorated stink bug (BMSB), corn earworm (CE), and spongy moth (SM) lose approximately 10–41 % of their suitable area, while the spotted lanternfly (SL) declines by about 42–46 % under most scenarios. In contrast, the root weevil (RW) expands, gaining 56–72 % more suitable habitat. By 2060, trends diverge more clearly across climate pathways. Under the low–emissions scenario SSP1–2.6, habitat loss remains moderate: suitable area decreases by approximately 28 % for BMSB, 39 % for CE, 32 % for SM, and 38 % for SL. RW continues to expand, gaining about 69 %.

Under intermediate warming (SSP2–4.5), declines intensify. Suitable habitat shrinks by 10 % for BMSB, 66 % for CE, 39 % for SM, and 66 % for SL, while RW suitable area increases by nearly 79 %. For SSP3–7.0, losses become more severe. Habitat decreases by approximately 35 % for BMSB, 59 % for CE, 44 % for SM, and 61 % for SL. RW suitable area continues to expand, increasing by roughly 88 %. Under the high–emissions scenario SSP5–8.5, habitat declines further for most species: approximately 39 % for BMSB, 41 % for CE, 47 % for SM, and 52 % for SL. RW remains the only species to gain habitat, increasing by about 58 %.

Model uncertainty also increases under stronger warming. For example, predicted habitat suitability for CE under SSP5–8.5 shows a standard deviation of approximately ±1.33 %, compared with only ±0.11 % under the current climate. This widening spread reflects greater variability across climate models and heightened sensitivity of species responses under intensified climate stress.

All insect species considered in this study exhibit a consistent response, with their climatically favorable habitats gradually shifting northward. This northward shift can be seen as a direct effect of global warming. As global temperatures rise, the *Goldilocks* zone for insects—where conditions are not too hot, not too cold, and sufficiently humid—tends to move northward Bebber et al. (2013); Skendžić et al. (2021); Schattman et al. (2024); Osland et al. (2021).

Previous modeling efforts for these insect species also reveal a consistent pattern of northward range expansion. In table 8, we summarize studies that projected habitat suitability in the future for the brown marmorated stink bug, corn earworm, and spotted lanternfly. Our findings strongly align with these studies in identifying a northward migration trend. However, we observe some discrepancies in the patterns of habitat contraction and expansion.

**Table 8:** Summary of previous modeling and observational studies on habitat suitability shifts for pest insects under climate change.

| Species | Model Type / Method | Geographical Extent | Northward Shift | Key Findings / Variables |
| --- | --- | --- | --- | --- |
| BMSB | CLIMEX (Kistner et al. Kistner (2017)) | Global | Yes | Northward expansion; contraction in southern US due to heat stress. Incorporated physiological thresholds (e.g., temperature, moisture, stress indices, diapause). |
|  | MaxEnt + BRT (Illán et al. Illán et al. (2022)) | Contiguous US | Yes | Increase in suitable habitat. Key predictors: January/summer max temperature and precipitation, growing degree day (GDD), distance to urban areas, land cover, elevation. |
| CE | Process-based (Diffenbaugh et al. Diffenbaugh et al. (2008)) | Contiguous US | Yes | Used biological thresholds (e.g., minimum winter temp, GDD). Projected strong northward expansion under future climate scenarios. |
| SL | MaxEnt (Zhao et al. Zhao et al. (2024)) | Global | Yes | Used 19 bioclimatic variables + elevation; BIO9 (mean temperature of driest quarter) was most influential. Projected global northward expansion; limited details for US. |

While no SDM-based projections exist for spongy moth and root weevil, observational evidence suggests similar trends. Spongy moth has shown a northward expansion into Ontario, Canada, with new infestations in previously unaffected areas Régnière et al. (2009); Srivastava et al. (2020). For root weevil, the eggs are highly cold-sensitive, dying after four days at 12°C (53°F) Lapointe et al. (2014). This constraint currently limits the species to the southern US, though warming winters may enable future expansion on Plant Health (PLH).

Given that MaxEnt predictions are highly sensitive to the geographical extent, the specific environmental variables used, and model settings, we suggest caution before reaching any strong conclusions while comparing different studiesMerow et al. (2013); Warren and Seifert (2011); Phillips and Dudík (2008); Amaro et al. (2023); VanDerWal et al. (2009); Low et al. (2021); Moreno-Amat et al. (2015).

### 3.3. Feature importance

In this study, we examine not only how habitat suitability changes under different climate scenarios, but also which climatic factors are most responsible for these changes. To do this, we analyze permutation importance, a widely used approach for identifying the features that most influence MaxEnt model predictions. Table 9 summarizes the top five contributing variables across 20 model runs for the brown marmorated stink bug, including how frequently each variable was selected and its average importance score. All runs used the same model settings and occurrence data, with variation introduced only through different random seeds for background sampling. Although the experimental settings remained largely consistent, we observed noticeable variation in both the identity of the most influential variables and their associated importance scores.

**Table 9:** Top 5 important features with their permutation score for brown marmorated stink bug under SSP2-4.5 climate scenario.

| $\beta_j$ | permutation importance<br>ranking | variable, frequency, score |
| --- | --- | --- |
| 0.5 | 1 | (Bio4, 5, 0.21); (elevation, 5, 0.182); (Bio2, 10, 0.196) |
| 0.5 | 2 | (Bio7, 5, 0.188); (Bio2, 5, 0.148); (elevation, 10, 0.193) |
| 0.5 | 3 | (elevation, 5, 0.187); (Bio4, 15, 0.12) |
| 0.5 | 4 | (Bio14, 15, 0.102); (Bio3, 5, 0.092) |
| 0.5 | 5 | (Bio13, 6, 0.057); (Bio3, 10, 0.084); (Bio14, 4, 0.074) |
| 1.0 | 1 | (elevation, 20, 0.181) |
| 1.0 | 2 | (Bio14, 5, 0.125); (Bio2, 5, 0.15); (Bio4, 5, 0.094); (Bio7, 5, 0.17) |
| 1.0 | 3 | (Bio2, 4, 0.118); (Bio4, 12, 0.113); (Bio1, 4, 0.062) |
| 1.0 | 4 | (Bio4, 4, 0.082); (Bio14, 12, 0.083); (Bio2, 4, 0.061) |
| 1.0 | 5 | (Bio3, 11, 0.058); (Bio5, 5, 0.055); (Bio15, 4, 0.049) |
| 1.5 | 1 | (elevation, 20, 0.178) |
| 1.5 | 2 | (Bio2, 10, 0.101); (Bio4, 10, 0.078) |
| 1.5 | 3 | (Bio4, 11, 0.088); (Bio14, 4, 0.074); (Bio1, 5, 0.064) |
| 1.5 | 4 | (Bio3, 5, 0.055); (Bio2, 5, 0.071); (Bio14, 5, 0.056); (Bio5, 5, 0.057) |
| 1.5 | 5 | (Bio14, 5, 0.05); (Bio1, 5, 0.054); (Bio13, 5, 0.038); (Bio2, 5, 0.054) |
| 2.0 | 1 | (elevation, 20, 0.194) |
| 2.0 | 2 | (Bio7, 13, 0.146); (Bio4, 7, 0.074) |
| 2.0 | 3 | (Bio4, 13, 0.133); (Bio2, 7, 0.073) |
| 2.0 | 4 | (Bio6, 14, 0.039); (Bio14, 6, 0.056) |
| 2.0 | 5 | (Bio15, 6, 0.039); (Bio3, 9, 0.04); (Bio13, 5, 0.041) |
| 2.5 | 1 | (elevation, 20, 0.177) |
| 2.5 | 2 | (Bio4, 9, 0.128); (Bio7, 4, 0.138); (Bio2, 7, 0.103) |
| 2.5 | 3 | (Bio7, 10, 0.119); (Bio4, 10, 0.1) |
| 2.5 | 4 | (Bio14, 14, 0.042); (Bio3, 6, 0.051) |
| 2.5 | 5 | (Bio3, 8, 0.033); (Bio2, 6, 0.051); (Bio13, 6, 0.034) |

We observe certain variables act as consistent predictors of suitability, while others may be more sensitive to the configuration of background environments. For example, BIO1 (annual mean temperature) appeared among the top three predictors in 18 out of 20 runs with relatively low variance in importance scores, suggesting it is a robust environmental driver for brown marmorated stink bug. In contrast, variables like BIO14 (precipitation of the driest month) appeared less frequently and exhibited greater score variability, indicating that their importance may depend on specific background contrasts or regional interactions.

To better understand the variability observed in permutation importance, we performed some controlled experiments. Using a simplified setup for the brown marmorated stink bug and including only bioclimatic variables, we systematically introduced spatial noise to the background points. Specifically, we generated *drifted* background datasets by randomly shifting each background point by up to 0 (no drift), 1, 5, 10, or 50 kilometers from its original location. The results of this analysis are presented in fig. 11. Even with a minimal displacement of just 1 km, we observed a change in feature importance rankings. For example, Bio19, which was the fourth most important feature in the base case, became the top-ranked feature after the 1 km drift. While the specific order and magnitude change, certain bioclimatic variables (e.g., Bio2, Bio4, Bio15, Bio19, Bio5, Bio6), appear repeatedly across different ranks and drift levels.

**Figure 11:**
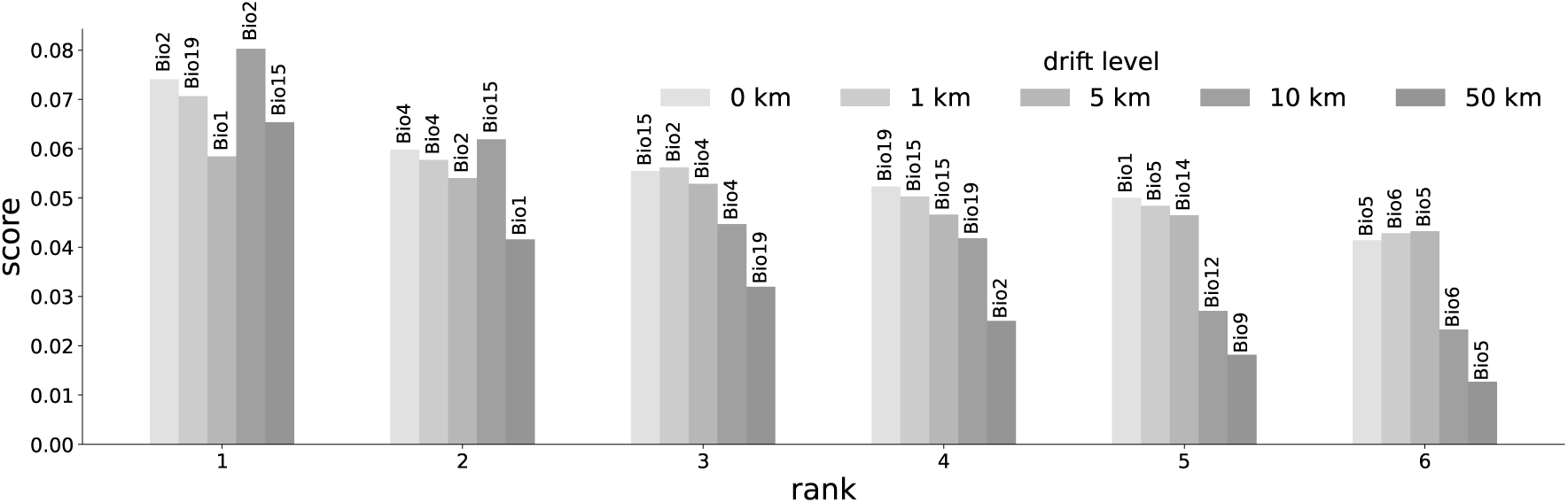
Comparison of top 6 important features for the drift dataset with their permutation score for the brown marmorated stink bug under SSP2-4.5 climate scenario.

Both analyses suggest that even modest changes in parameter settings can lead to shifts in the ranking of feature importance and their score. Therefore, relying on a strict feature ranking to draw ecological insights or inform decision-making should be approached with caution. We recommend that, instead of relying on a single experiment to assess feature importance, researchers should identify features that consistently appear among the top contributors across multiple runs. Features that appear less frequently can confidently be avoided.

## 4. Discussion

We study the current and future habitat of five important insect species over the contiguous US using MaxEnt. We looked into four different climate change scenarios, and investigated the influence of model settings on MaxEnt prediction. Our goal in this study was threefold: (1) to understand how the choice of background points influences MaxEnt’s predictions, (2) to evaluate how these predictions change under future climate scenarios, and (3) to assess the robustness of feature-importance rankings.

Our analysis of the background point distribution revealed that a modest buffer bias (mixing coefficient of *m* ≈ 0.25) consistently yielded the best CBI scores for most experiments. This suggests that using a hybrid approach rather than purely uniform sampling (*m* = 0) or highly localized sampling (*m* = 1) yields the best calibrated model. We recommend using *m* as a regularization parameter, but a default value of *m* = 0.25 can be a good starting point. We further demonstrated that the AUC, while a commonly used metric, remains a poor approach over CBI.

Our projections of suitable habitat under various future climate scenarios (SSP1–2.6, SSP2–4.5, SSP3–7.0, and SSP5–8.5) revealed inconsistent patterns of habitat expansion and contraction in the future. Our experiments demonstrate that these projections are highly sensitive to the settings used during model training. Certain configurations result in an expansion of suitable habitat, while others predict a reduction. As discussed in Section 3.2, many studies use the default settings of MaxEnt when modeling insect habitat suitability. Under default MaxEnt settings and a moderate emission scenario, we observe a general northward expansion of insect distributions. All insect species, except root weevil, show a reduction in future suitable habitat; root weevil, in contrast, shows an increase. We also emphasize that comparisons across studies should be made with caution, as variations in modeling settings, background sampling, and the geographical extent of the study can lead to fundamentally different interpretations of species’ future distributions.

Finally, our investigation into feature importance underscored the considerable variability in variable rankings and scores, even within a single experimental setting and with minor perturbations to background samples. We found that even a minimal spatial displacement of background points significantly altered the permutation importance rankings. While some variables, such as annual mean temperature (BIO1) for brown marmorated stink bug, consistently emerged as robust environmental drivers, others exhibited greater sensitivity to background contrasts or regional interactions. This calls into question the practice of relying on a single experiment to assess feature importance for ecological insights or decision-making. Instead, we advocate for identifying features that consistently appear among the top contributors across multiple runs, thereby increasing confidence in their ecological relevance.

In conclusion, this study provides crucial insights into the complexities of species distribution modeling using MaxEnt. We demonstrate that careful consideration of background sampling strategies, rigorous evaluation using calibration-focused metrics, and thorough sensitivity analyses of both future habitat projections and feature importance are paramount for generating robust and reliable predictions. Our findings serve as a cautionary reminder against over-reliance on default settings and emphasize the need for a more comprehensive and critical approach to MaxEnt model application in ecological research and conservation planning.

## Supporting information

SI information for manuscript

## 5. Acknowledgements

This research was funded by the U.S. Department of Agriculture (USDA), National Institute of Food and Agriculture (NIFA), through Florida A&M University (Accession No. 1028629, Project Director: Muhammad Haseeb).

During the preparation of this work the author(s) used ChatGPT to edit the language and grammar of the manuscript. After using this tool/service, the author(s) reviewed and edited the content as needed and take(s) full responsibility for the content of the publication.

