## Supplementary material for "Evaluating MaxEnt Modeling Strategies for Predicting Suitable Habitats of Invasive Insects Under Climate Change Scenarios": SI information for manuscript

---

---

### Contents

**1 Insects Considered 2**

### 1. Insects Considered

We considered five key pest species in this study, which posea threat to US agriculture. Here we are summarizing their native range, host preferences, and temperature–humidity tolerances.

- Brown marmorated stink bug (Halyomorpha halys): The brown marmorated stink bug is a pest native to Asia (China, Japan, Korea). It feeds on many plants but prefers fruiting plants like apples, pears, peaches, grapes, blueberries, soybeans, tomatoes, and corn. Though it mainly feeds on fruits, it can also damage other plant parts and even pierce tree bark [7]. This bug thrives in warm temperatures (25–30°C)[2, 6]. High temperatures and low humidity reduce its survival, especially in dry conditions. In California, high heat and low humidity have been linked to a decline in its population [2]. Low temperatures don't promote growth; instead, they slow down development and encourage the bug to enter dormancy (diapause) during winter. Milder temperatures in spring and summer help the bug grow and reproduce more actively, leading to larger populations.

- Corn earworm (Helicoverpa zea): The corn earworm (Helicoverpa zea) is a pest native to the Americas that primarily feeds on corn but also damages crops like cotton, tomatoes, and soybeans. It thrives in warm, humid environments and remains active throughout the year in tropical and subtropical regions. In the southern United States (below 40° latitude), it can survive year-round, while in higher latitudes (above 40° latitude), it is only active during the summer. Hot and moist conditions help the corn earworm develop and spread more quickly, making infestations more severe. On the other hand, the corn earworm struggles to survive in cold and dry environments. It cannot withstand harsh winters, which limits its presence in colder regions. In areas with higher latitudes and colder climates, the pest disappears when temperatures drop and only returns when the weather warms up again. Overall, the corn earworm thrives in hot and humid climates, especially in lower latitudes, but faces survival challenges in colder, drier regions at higher latitudes.

- Spongy moth (Lymantria dispar): The spongy moth, formerly known as the gypsy moth, is an invasive forest pest native to Europe and Asia. It feeds on the leaves of over 300 species of trees and shrubs, with a strong preference for oak, birch, willow, and aspen [1]. The larvae are the primary feeding stage and can cause extensive defoliation, weakening trees, and making them vulnerable to other stresses. Spongy moth eggs need to experience a cold winter; however, they cannot survive in temperatures below -20 degrees Celsius. Temperature plays a key role in the moth's development: optimal larval development occurs at moderate temperatures (20–25°C), while extreme heat or cold can reduce survival rates [3].

- Root weevil (Diaprepes abbreviatus): Root weevils (Curculionidae family) are a diverse group of beetles found in various regions, with different species adapted to different climates and crops. For example, the Diaprepes root weevil (Diaprepes abbreviatus) is native to the Caribbean and has become a major pest in Florida and California, while the strawberry root weevil (Otiorhynchus ovatus) is widespread in Canada and the northern United States. These pests feed on a wide range of plants—Diaprepes targets over 270 species, including citrus, sugarcane, and ornamentals, while others focus on strawberries and shrubs [5]. Root weevils thrive in mild to warm temperatures (15–27°C), with warmer conditions accelerating their life cycles. Moist, poorly drained soils and high humidity provide ideal environments for larvae, especially in greenhouses. In contrast, cold winters can help reduce populations, although weevils can survive in protected microhabitats [5].

- Spotted lanternfly (Lycorma delicatula): The spotted lanternfly is native to China, India, and Vietnam, and has become a serious invasive pest in the United States. This insect is highly polyphagous, feeding on more than 70 plant species. Its preferred hosts include grapevines (Vitis spp.), apples (Malus domestica), stone fruits (e.g., peaches, plums, cherries), and hardwood trees. The spotted lanternfly develops best at temperatures between 15–30°C, with faster growth in warmer conditions [8]. Although less studied, high temperatures combined with low humidity may increase mortality, suggesting an interaction between temperature and humidity in population dynamics [4].
